# Genome-wide association and genomic prediction of growth traits in the European flat oyster (*Ostrea edulis*)

**DOI:** 10.1101/2022.06.10.495672

**Authors:** Carolina Peñaloza, Agustin Barria, Athina Papadopoulou, Chantelle Hooper, Joanne Preston, Matthew Green, Luke Helmer, Jacob Kean Hammerson, Jennifer Nascimento-Schulze, Diana Minardi, Manu Kumar Gundappa, Daniel J Macqueen, John Hamilton, Ross D Houston, Tim P Bean

## Abstract

The European flat oyster (*Ostrea edulis*) is a bivalve mollusc that was once widely distributed in Europe and represented an important food resource for humans for centuries. Populations of *O. edulis* experienced a severe decline across their biogeographic range mainly due to anthropogenic activities and disease outbreaks. To restore the economic and ecological benefits of European flat oyster populations, extensive protection and restoration efforts are in place within Europe. In line with the increasing interest in supporting restoration and oyster farming through the breeding of stocks with enhanced performance, the present study aimed to evaluate the potential of genomic selection for improving growth traits in a European flat oyster population obtained from successive mass-spawning events. Four growth-related traits were evaluated: total weight (TW), shell height (SH), shell width (SW) and shell length (SL). The heritability of the growth traits was moderate-low, with estimates of 0.45, 0.37, 0.22, and 0.32 for TW, SH, SW and SL, respectively. A genome-wide association analysis revealed a largely polygenic genetic architecture for the four growth traits, with two distinct QTLs detected on chromosome 4. To investigate whether genomic selection can be implemented in flat oyster breeding at a reduced cost, the utility of low-density SNP panels (down to 100 SNPs) was assessed. Genomic prediction accuracies using the full density panel were high (>0.83 for all traits). The evaluation of the effect of reducing the number of markers used to predict genomic breeding values revealed that similar selection accuracies could be achieved for all traits with 2K SNPs as for a full panel containing 4,577 SNPs. Only slight reductions in accuracies were observed at the lowest SNP density tested (i.e. 100 SNPs), likely due to a high relatedness between individuals being included in the training and validation sets during cross-validation. Overall, our results suggest that the genetic improvement of growth traits in oysters is feasible. Nevertheless, and although low-density SNP panels appear as a promising strategy for applying GS at a reduced cost, additional populations with different degrees of genetic relationship should be assessed to derive estimates of prediction accuracies to be expected in practical breeding programmes.

## 1. INTRODUCTION

The European flat oyster (*Ostrea edulis*) was an abundant native bivalve species and an important fishery resource in much of Europe up to the 19^th^ century (Pogoda 2019). However, populations of *O. edulis* experienced a severe decline across their biogeographic range due to a range of detrimental factors including overfishing and habitat degradation (Thurstan et al. 2013), the subsequent invasion of non-native species (e.g. slipper limpet, *Crepidula fornicata*) (Preston et al. 2020; Helmer et al. 2019) and pathogenic diseases (Robert et al. 1991; Sas et al. 2020). The continuous decimation of native populations in the Atlantic and Mediterranean seas led to significant changes in oyster production, which progressively shifted towards farming (Korringa 1976), and eventually to the cultivation of different species including *Crassostrea angulata* (Oelig and Uf 2000) and the non-indigenous Pacific oyster (*Crassostrea gigas*) (Grizel and Héral 1991; Walne and Helm 1979). The Pacific oyster was introduced into Europe for aquaculture purposes owing to its favourable production traits, such as a faster growth rate and higher resistance to the main diseases affecting *C. angulata* and *O. edulis* (Renault et al. 1995; Grizel and Héral 1991). Worldwide oyster production is now dominated by the Pacific oyster (97.7%), while the production of the European flat oyster remains stably low, constituting just ∼0.2% of global production (FAO 2019). Despite the demand for shellfish continues to increase (Botta et al. 2020), the level of *O. edulis* production is stagnant. One of the main factors that hinders the growth of the industry is the lack of a substantial and steady supply of oyster seed (i.e. juveniles) (see Colsoul et al. (2021) for a review). Hence, the optimization of oyster larval production in hatcheries and spatting ponds is key for future European flat oyster aquaculture, as well as for restoration projects, which are also expected to rely on sustainable sources of juveniles for restocking (Pogoda et al. 2020). Importantly, the artificial propagation of flat oyster seed will facilitate the application of selective breeding programmes. Although selective breeding programmes are typically used to improve aquaculture production, they could also benefit the ecological restoration of *O. edulis*. If desirable traits such as disease resistance are found to have a strong genetic component, then increased resistance to life-limiting diseases – such as bonamiosis (Culloty et al. 2004; Naciri-Graven et al. 1998) – could potentially be achieved while maintaining the adaptive potential (i.e. genetic diversity) of restored populations.

Selective breeding in oysters has mainly focused on improving meat yield and quality, disease resistance, survival and growth (Toro and Newkirk, 1990; Allen et al., 1993; Ragone Calvo et al., 2003; Ward et al., 2005; Dégremont et al., 2015; De Melo et al., 2016; Proestou et al., 2016; Camara et al., 2017; Zhang et al., 2019), with a recent interest in nutritional content and shell shape (Grizzle et al., 2017; Liu et al., 2019; Meng et al., 2019; Wan et al., 2020; He et al., 2022). Among these traits, growth is comparatively simple to assess and consequently select for using phenotypic information. Although the direct comparison of heritability estimates from different studies is difficult (e.g. due to intrinsic differences between populations), estimates for growth rate in oysters tend to be moderate (e.g. 0.26 and 0.31 – De Melo et al. (2016) and Evans and Langdon (2006), respectively). As a result, fast-growing lines of oysters have been developed for some of the main commercial species, such as the Pacific (*C. gigas*) (Zhang et al. 2019), Portuguese (*C. angulata*) (Vu et al. 2020), American (*C. virginica*) (Varney and Wilbur 2020) and Sydney rock (*Saccostrea glomerata*) (Fitzer et al. 2019) oyster. Initial attempts to genetically improve the European flat oyster *O. edulis* resulted in a 23% increase in growth rate compared to an unselected (control) line (Newkirk and Haley 1982). This striking genetic response was not replicated in a second generation of selection, possibly due to unintentional inbreeding (Newkirk and Haley 1983). Indeed, even relatively modest levels of inbreeding have been shown to significantly affect performance traits in oysters (Evans et al. 2004), highlighting the importance of an adequate management of genetic diversity in hatchery-derived stocks. Moreover, oysters and bivalves in general, appear to have a high genetic load (see for a review Plough (2016)) and, therefore, may be particularly susceptible to inbreeding depression. Hence, the incorporation of genomic tools into shellfish breeding schemes will be key for balancing genetic gain with population diversity in order to sustain the long-term progress for traits under selection.

A vast array of genomic tools and resources have become available for genetic research and breeding applications in oysters. For example, for economically relevant species, chromosome-level genome assemblies (Peñaloza et al. 2021; Qi et al. 2021; Modak et al. 2021; Li et al. 2021), SNP arrays (Gutierrez et al. 2017; Qi et al. 2017; Lapegue et al. 2014) and medium-density linkage maps (Gutierrez et al. 2018; Li et al. 2018; Jones et al. 2013; Wang et al. 2016; Yin et al. 2020) have been produced. These resources have been applied to examine the genetic basis of growth (Gutierrez et al. 2018; Jones et al. 2014; He et al. 2021), low salinity tolerance (McCarty et al. 2021), disease resistance (Gutierrez et al. 2018; Yang et al. 2022) and nutritional content (Meng et al. 2019). For the European flat oyster, high-quality genomes have recently been released (Boutet et al. 2022; Gundappa et al. 2022), which along with available high-throughput genotyping techniques (e.g. SNP arrays and genotype-by-sequencing approaches), provide the opportunity for gaining insight into the genomic architecture of relevant production traits. Most of the traits of economic importance in aquaculture species have a polygenic architecture (Zenger et al. 2019). For polygenic traits (i.e. those controlled by many loci), the application of predictive techniques such as genomic selection (GS) may enable a faster genetic gain than conventional pedigree-based selection. GS is a method based on genome-wide markers in which the effect of all loci are simultaneously used for predicting the estimated breeding values (EBV) of selection candidates (Meuwissen et al. 2001), and has shown major potential in aquaculture species, where it can be used to characterise variation within and between large families of potential breeders. However, commercial application to aquaculture production is largely limited to the major finfish and crustacean species (e.g. salmonids, Nile tilapia, tropical shrimp) (Lillehammer et al. 2020; Boudry et al. 2021; Zenger et al. 2019). Studies into the feasibility of applying genomic selection schemes in oyster breeding programmes have shown that for growth (Vu et al. 2021b; Gutierrez et al. 2018), edibility (Vu et al. 2021b), low salinity tolerance (McCarty et al. 2021), and disease resistance traits (Vu et al. 2021b; Gutierrez et al. 2020), greater genetic gains could be achieved through GS compared to traditional breeding. Nevertheless, the practical application of GS as a selection strategy will likely depend on how cost-effective it is compared to pedigree-based methods. The development of feasible alternatives for reducing genotyping costs, such as using affordable low-density genotyping tools that yield similar accuracies than higher-density panels, will be critical for the potential of GS to be realized by oyster breeding programmes.

In line with the increasing interest in supporting oyster culture and restoration through the breeding of stocks with enhanced performance, the overall aim of this study was to evaluate the potential of GS for the genetic improvement of growth and growth-related (morphometric) traits in the European flat oyster. First, the heritability of total weight, shell length, shell width and shell height was estimated for a hatchery-derived population genotyped using a ∼15K SNP array. Second, a genome-wide association (GWAS) analysis was conducted to dissect the genetic architecture of the measured traits. Last, to evaluate whether GS may be an effective and cost-effective strategy for improving traits associated with oyster growth, the accuracy of genomic predictions using reduced density SNP marker panels was assessed.

## 2. MATERIALS AND METHODS

### 2.1 Field experiment

The European flat oyster population used in this study was generated in a UK-based hatchery (Sea Salter Morecombe hatchery) by mass spawning of approximately 40 broodstock parents over several spawning events. The resulting F1 generation was then deployed to Lochnell oysters (56.494° N, 5.459° W) and grown for six months in ortac grow-out cages. Animals were then transferred to the Institute of Marine Sciences at the University of Portsmouth (UK), and maintained in a flow-through system until deployment. During this holding period, ∼1,000 randomly selected oysters were individually tagged and their first phenotype measurements recorded (see ‘Phenotypes’ section below). Prior to deployment, animals were cleaned of fouling, washed in fresh water and dried. Embossed plastic tags with unique identifier codes were attached to their shell with epoxy resin glue. Animals were returned to aquaria within the hour. Oysters were placed in Aquamesh® cages (L 0.55 m x W 0.55 m x D 0.4 m) (GT Products Europe Ltd) at a density of 200 oysters per cage, and deployed 1 metre below floating pontoons at Port Hamble Marina (MDL) in the River Hamble (50.861° N, 1.312° W) in January 2019. Mortalities were documented monthly and dead oysters – i.e. those with empty or gaping shells – were removed from the experiment. General disease status was assessed on subsets of oysters throughout the experiment by histology and *in situ* hybridisation using an adaptation of available methods (Fabioux et al. 2004; Montagnani et al. 2001). In addition, the presence of *Bonamia ostreae*, a protozoan parasite that causes a lethal infection of flat oyster haemocytes (Pichot et al. 1979), was assessed by qPCR following Robert et al. (2009). The prevalence of *B. ostreae* infections was negligible; hence disease status had a minor influence on the assessment of growth traits in the experimental population. After 10 months of growth under field conditions, gill tissue was dissected from individuals alive at the end of the study and preserved in molecular grade absolute ethanol (Fisher Scientific) for genetic analysis.

### 2.2 Phenotypes

Four growth-associated traits were measured at three time points over the course of 10 months: total weight (TW, the weight of an individual oyster including the shell), shell length (SL, the maximum distance between the anterior and posterior margins), shell height (SH, the maximum distance between the hinge to the furthermost edge), and shell width (SW, the maximum distance at the thickest part of the two shell valves) (Figure 1). Weight was recorded in grams up to one decimal place. Shell measurements were taken with traceable digital callipers (Fisher Scientific) with 0.02 mm precision. Oysters were cleaned and defouled before measurements were taken.

**Figure 1.**
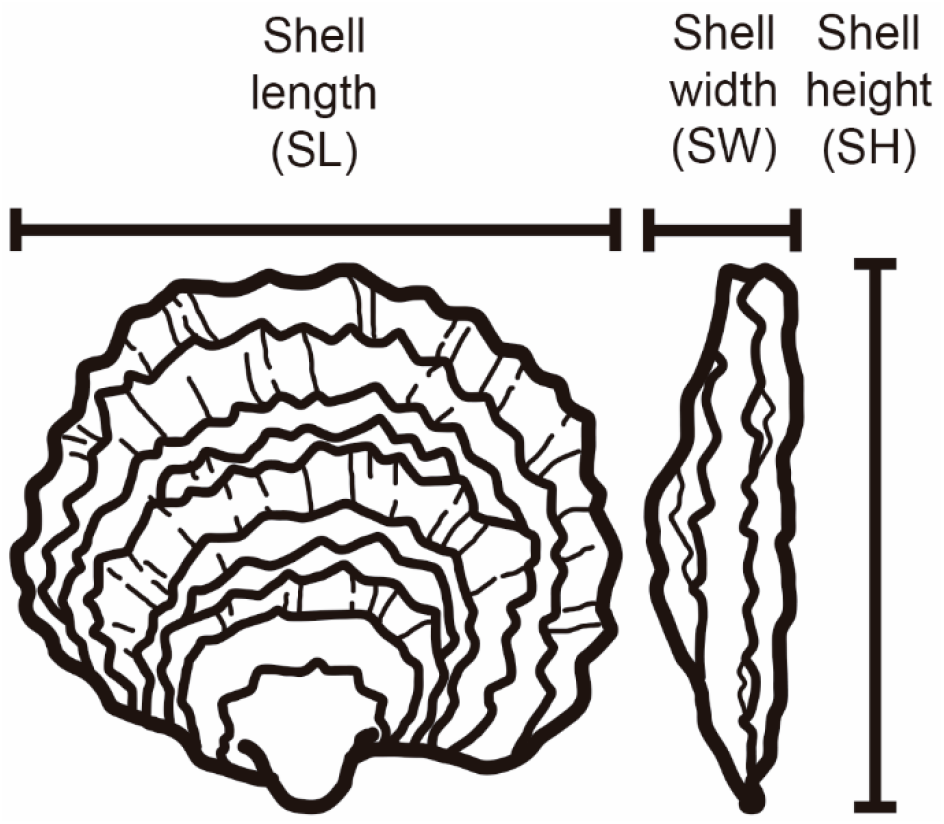
Nomenclature of the growth-related morphometric traits measured in this study.

### 2.3 DNA extraction

Total DNA was isolated from gill tissue following a CTAB (cetyltrimethylammonium bromide)- based extraction protocol (details in Gutierrez et al. (2017)). The integrity of the extracted DNA was assessed by agarose gel electrophoresis, while DNA quality was verified on a Nanodrop ND-1000 (Thermo Fisher Scientific) spectrophotometer by checking the 260/280 and 260/230 ratios. All samples had 260/280 and 260/230 values ≥1.85 and ≥1.96, respectively.

### 2.4 SNP genotyping and Quality Control

Whole-genome genotyping of ∼15K SNPs was carried out by IdentiGEN (Dublin, Ireland) using the combined-species Affymetrix Axiom oyster SNP-array (Gutierrez et al. 2017). Signal intensity files were imported to the Axiom analysis Suite v4.0.3.3 software for quality control (QC) assessment and genotype calling. Genotypes were generated using the default parameter settings for diploid species, resulting in 11,808 SNPs typed for 870 individuals. To assess the reproducibility of genotype calls, five DNA samples from the same individual were genotyped independently on three different arrays, and their genotype concordance evaluated through an identity-by-state (IBS) analysis. The genotype concordance rate among replicates was 99.7%, demonstrating a high reproducibility of the genotyping assays. The flanking region of these markers were mapped to the *O. edulis* chromosome-level genome assembly (Gundappa et al. 2022). Of the 11,808 SNPs, 10,025 had uniquely mapping probes and were retained for downstream analysis. A total of 1,539 markers (15.4%) were monomorphic in the population under study. QC was conducted using Plink v2.0 (Chang et al. 2015). SNP variants were retained for further analysis if they had a call rate >95% and a minor allele frequency (MAF) >0.05. These filters removed 4,391 SNPs (leaving a total of 5,634 SNPs), of which the majority were filtered out based on the MAF threshold (i.e. were monomorphic or near-monomorphic in this population). Given that significant sub-clustering was detected in the data (Figure S2), possibly due to a high variance in the reproductive success of broodstock parents and/or temporal variation in spawning, a k-means clustering method was used to assign individuals into groups. Deviations from Hardy-Weinberg Equilibrium (HWE) were tested separately in each of the three genetic clusters identified by the analysis. SNP markers showing significant deviations (HWE p-value < 1e-10) in two of the three clusters were excluded from the analysis. Sample QC included removing individual oysters with a missingness above 5% and high heterozygosity (i.e. more than three median absolute deviations from median). Finally, a principal component analysis (PCA) was performed using a set of ∼3.5K SNPs for which no pair of markers within a window of 200 kb had a *r*^2^ >0.5. The top five PCs, which explain 47% of the variance (considering 20 PCs), were fitted in the model to account for the effect of population structure. The final dataset comprised 840 samples genotyped at 4,577 genome-wide SNPs.

### 2.5 Genetic parameter estimation

Genetic parameters for growth-related traits were estimated by fitting the following univariate linear mixed model in GEMMA v0.95alpha (Zhou and Stephens 2012):

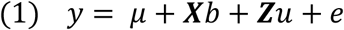

Where ***y*** is the vector of observed phenotypes; **μ** is the overall mean of the phenotype in the population; **b** is the vector of fixed effects to be fitted (the first five principal components were included as covariates); **u** is the vector of the additive genetic effects; ***X*** and ***Z*** are the corresponding incidence matrices for fixed and additive effects, respectively; and *e* is a vector of residuals. The following distributions were assumed: 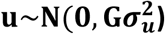 and 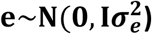. Where 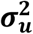 and 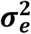 are the additive genetic and residual variance, respectively, **G** is the genomic relationship matrix and **I** is the identity matrix. The heritability of growth-related traits was estimated as the ratio of the additive genetic variance to the total phenotypic variance.

Bivariate animal linear models were implemented to estimate the genetic (co)variance between TW, SL, SH and SW. Each bivariate analysis was fitted with the same top 5 PCs mentioned above. Subsequently, genetic correlations among traits were measured as the ratio of the covariance of two traits to the square root of the product of the variance for each trait. Phenotypic correlations between traits were calculated using the Pearson correlation coefficient.

### 2.6 Genome-wide association study (GWAS)

To identify SNPs in the flat oyster genome correlated with variation in growth-related traits, a GWAS was performed by implementing the same model described previously in the GEMMA software. SNPs were considered significant at the genome-wide level if their likelihood ratio test P-values surpassed a conservative Bonferroni-corrected significance threshold (α/4,577 = 1.09e-5). To derive a threshold for chromosome-wide (suggestive) significance, α was divided by the average number of SNPs per chromosome (α/457 = 1.09e-4). The single-marker P-values obtained from GEMMA were plotted against their chromosome location using the R package qqman v 0.1.4 (Turner 2017). To assess the inflation of the association statistics, the genomic control coefficient lambda λGC was calculated following (Devlin and Roeder 1999). Candidate genes were searched within 100 kb of the most significant SNP loci using BEDOPS v2.4.26 (Neph et al. 2012).

### 2.7 Genomic Prediction

To evaluate the accuracy of genomic selection, a 5-fold cross validation approach - animals split into training (80%) and validation (20%) sets - was used on a population of 840 oysters genotyped for 4,577 informative SNP markers. To reduce stochastic effects arising from individual sampling, each analysis was repeated 10 times. For each replicate, animals were randomly partitioned into five subsets (each subset contained 168 individuals). TW, SL, SH and SW phenotypes recorded in individuals allocated to one of the subsets (validation set) were masked. The breeding values of the validation set were then predicted based on the information from the remaining four subsets (training sets) using model (1). The model was fitted using the AIREMLF90 module from BLUPF90. The accuracy of genomic predictions was calculated as follows:

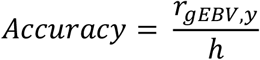

where r_gEBV,y_ is the correlation between the predicted and the actual phenotypes of the validation set, while *h* is the square root of the heritability of the trait estimated as described above.

### 2.8 Evaluation of the effect of SNP density on genomic predictions

To assess the effect of SNP density on the accuracy of genomic predictions of growth-related traits, SNP panels of varying sizes were randomly sampled from the final pool of QC-filtered array markers (n = 4,577 SNPs). Panels of the following densities were evaluated: 4K, 3K, 2K, 1K, 500, 400, 300, 200 and 100 SNPs. To build the lower-density panels, markers were randomly sampled from the full QC-filtered SNP dataset in proportion to chromosome lengths using the R package CVrepGPAcalc v1.0 (Tsairidou 2019; Tsairidou et al. 2020). To account for sampling bias, 10 SNP panels were generated for each of the SNP densities. The average genomic prediction accuracies of the different low-density panels were compared against the equivalent accuracy values obtained with the full panel.

### 2.9 Data Availability

The phenotype data used in the current study can be found in Mendeley Data, https://doi.org/10.17632/sdtjyys7gr.1.

## 3. Results and discussion

### 3.1 Growth traits and heritability

Improvement of growth rate is typically one of the first traits to be included as a selection target in breeding programmes across many farmed species. In this study, oyster growth rate was assessed in a hatchery-derived oyster population that was translocated to a growing site and monitored for 10 months. The experimental population had a lower genome-wide heterozygosity (Ho=0.27; He=0.22) compared to the values reported by (Vera et al., 2019) (Ho>0.31) for a diverse set of flat oyster populations genotyped with the same array. An overall mortality of 14% was observed during the field trial, among which the majority (36%) occurred during a summer month (July). At the end of the experimental period, the *O. edulis* population had the following growth means and standard deviations: +15.7 g (SD = 5.8), 50.8 mm (SD = 7.3), 12.9 mm (SD = 2.3) and 45.8 mm (SD = 8.9), for TW, SH, SW, SL, respectively (Table 1). The phenotypic correlation was found to be the highest (*r* > 0.8) between two pairs of traits: (i) TW and SH, and (ii) TW and SL (Figure 2).

**Table 1.** Summary statistics of the phenotypic data (SD: Standard deviation; CV: coefficient of variation).

| Trait | Unit | Mean | Min | Max | SD | CV (%) |
| --- | --- | --- | --- | --- | --- | --- |
| Total weight | g | 15.7 | 4.0 | 38.5 | 5.8 | 36.8 |
| Shell height | mm | 50.8 | 22.9 | 76.1 | 7.3 | 14.4 |
| Shell width | mm | 12.9 | 6.5 | 27.7 | 2.3 | 18.2 |
| Shell length | mm | 45.8 | 22.2 | 94.4 | 8.9 | 19.5 |

**Figure 2.**
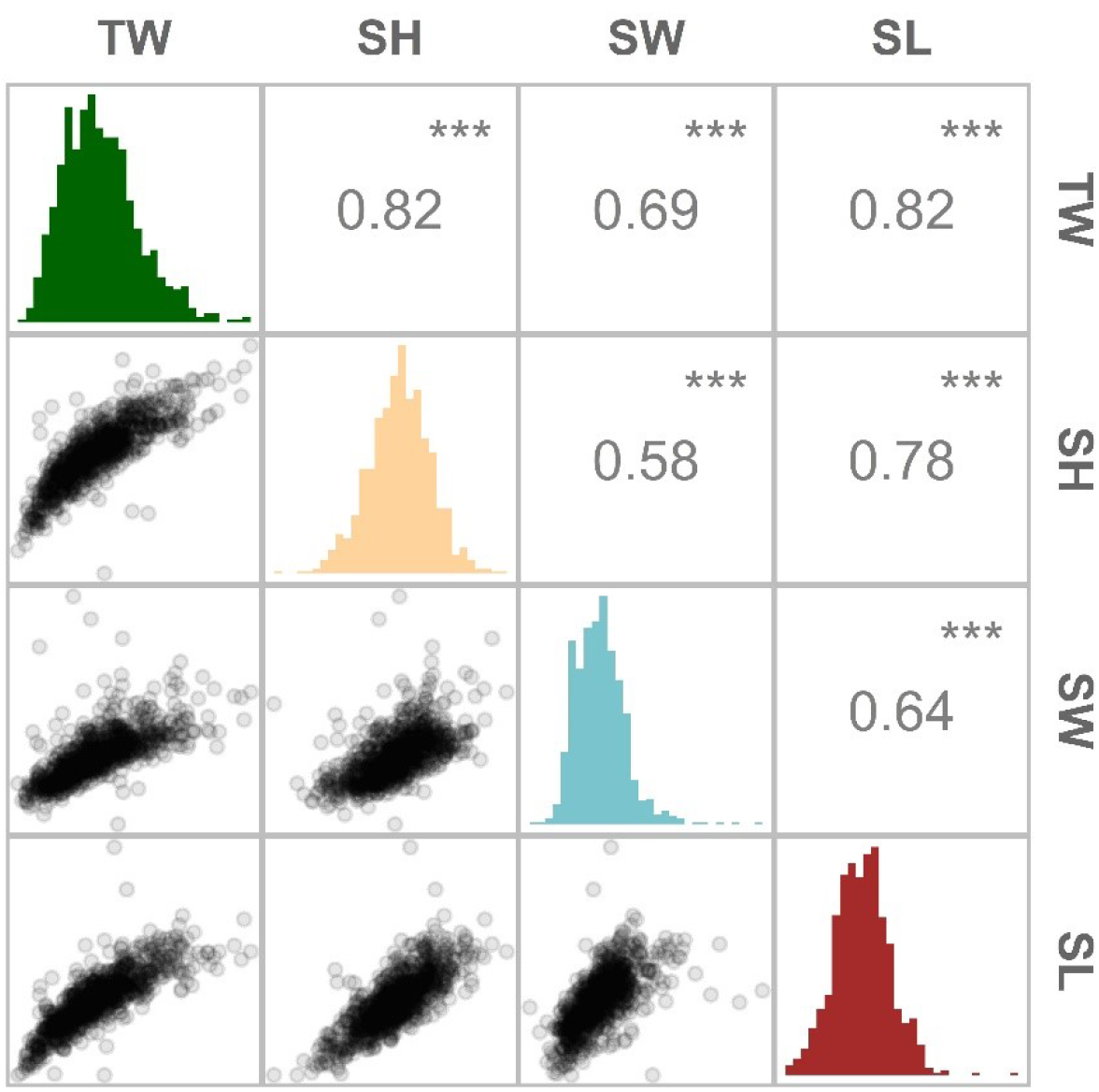
Distribution and magnitude of the phenotypic correlations between growth-related traits in *Ostrea edulis*. Pearson’s correlation between traits (above the diagonal), histogram of trait distribution (diagonal) and scatterplots comparing two traits (below the diagonal). TW (total weight), SH (shell height), SW (shell width) and SL (shell length). *** indicates *p*-values <0.001.

For the European oyster population under study, the heritability estimates of these growth-related traits were in the moderate range of 0.22 (for SW) to 0.45 (for TW) (Table 2). Consistent with similar studies carried out in related oyster species (Xu et al. 2017; Vu et al. 2020), heritability estimates based on SNP markers were higher for total weight than for growth-related morphometric traits (i.e. shell height, shell width and shell length). The estimation of heritability for total weight (herein referred to as TW) was similar to those reported for nine-month-old Portuguese oysters (*h*^2^ = 0.45) and a two-year old Pacific oyster strain (*h*^2^ = 0.42) (Vu et al. 2021b; Xu et al. 2017). Total weight, as measured in this study, is a composite phenotype made up of the animal’s shell and soft tissue weights, in addition to the weight of any pallial fluid - thus is not a direct reflection of meat yield. Nevertheless, in *C. angulata*, a positive genetic correlation (0.63) has been found between TW and soft tissue weight (Vu et al. 2021b), suggesting that selecting for TW - a trait easier to measure - could lead to improvements in meat yields. Such indirect improvements of correlated traits have been reported in a Portuguese oyster line selected only for harvest weight. While the achieved average selection response for total weight at harvest was 5.8% per generation, genetic gains were also observed for soft tissue weight, with indirect gains reaching a 1.2% increase per generation (Vu et al. 2020). For the shell-related traits examined in this study (SH, SW and SL), heritability estimates were in line with previous studies (Gutierrez et al. 2018; Yuehuan et al. 2017), and ranged from 0.22 to 0.37. Traditionally, the focus on shell morphometric traits was to improve oyster growth. Nevertheless, in recent years, oyster shell shape is increasingly being viewed as an attractive goal for selective breeding due to its growing importance for consumers (Mizuta and Wikfors 2019). The perceived attractiveness of an oyster shell can be represented as a secondary trait derived from a ratio between primary (shell dimension) traits, such as the shell width index (Kube et al. 2011). Given that significant heritable variation was observed for the three examined morphometric traits, strategies for homogenizing particular shell shapes may be feasible in *O. edulis*.

**Table 2.** Estimates of heritability (*h*^2^) and standard error on the diagonal and pairwise genetic correlations (below the diagonal) for growth-related traits in a European flat oyster population.

| Trait | Total weight | Shell height | Shell width | Shell length |
| --- | --- | --- | --- | --- |
| Total weight | **0.45 (0.06)** |  |  |  |
| Shell height | 0.99 | **0.37 (0.06)** |  |  |
| Shell width | 0.96 | 0.90 | **0.22 (0.05)** |  |
| Shell length | 0.95 | 0.93 | 0.88 | **0.32 (0.06)** |

### 3.2 Genome-wide association analysis for growth-related traits

A GWAS of ∼4.5K SNPs passing the filtering criteria were genotyped on 840 oysters with phenotypic records to gain insight into the genetic basis of growth rate variation in *O. edulis*. Three of the four examined traits showed association signals surpassing the genome-wide level of significance (Figure 3). The genomic inflation factor lambda of the GWAS analysis were close to the desired value (λ=1) (see Figure S1), indicating that population structure was adequately accounted for by the model. For TW, the GWAS identified two putative quantitative trait loci (QTLs) on chromosome 4 associated with the trait. The presence of two separate QTLs is supported by the low linkage disequilibrium observed between the most significant SNPs at each locus (pairwise *r*^2^ < 0.1). An additional 13 suggestive loci were also identified, of which nine were located in the vicinity of the two abovementioned genome-wide hits and four were found on chromosome 1 (Table S1). The SNP showing the strongest association with TW (AX-169174635) explained 3% of the phenotypic variance. This lead SNP was also found to be significantly associated with SH and SW. For SL, no SNP reached a genome-wide significance level, although a few of the same markers showing associations with TW, SH and SW surpassed the threshold for suggestive significance. The complete overlap of GWAS hits across the different traits suggests a high degree of shared genetic control among them, consistent with the high positive genetic correlations observed (Table 2). Overall, the GWAS results indicate that growth-related traits in *O. edulis* are influenced by many small-effect loci, exhibiting a polygenic architecture, but that two regions on chromosome 4 may have a moderate effect on these traits.

**Figure 3.**
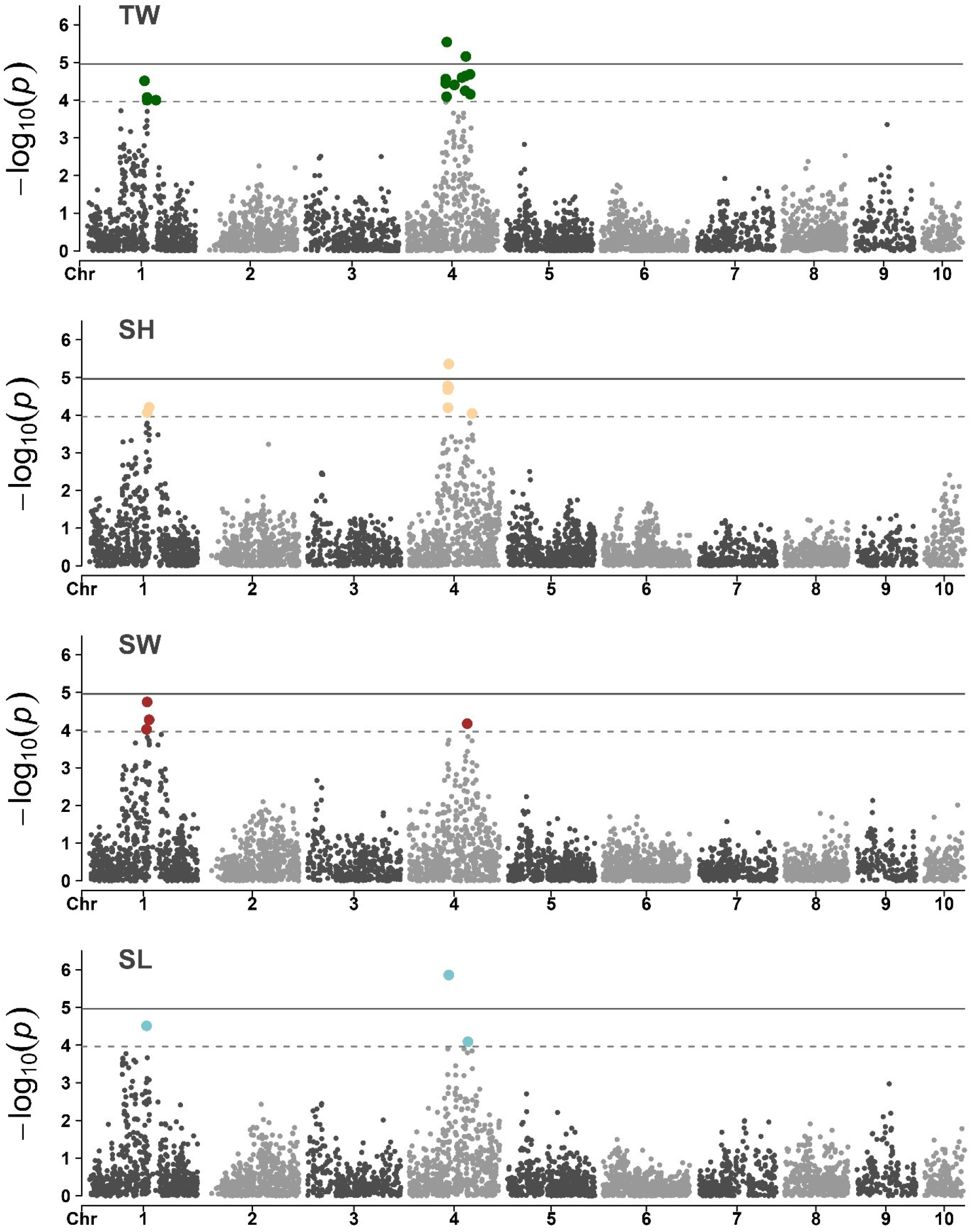
Manhattan plots of the GWAS for growth-related traits in a European flat oyster population. Solid lines indicates the threshold value for genome-wide significance. Dashed lines indicate the threshold for a suggestive (chromosome-level) significance.

The marker showing the most significant association with TW, SH and SW is located in the exon of a gene annotated as a *N4BP2* (NEDD4 Binding Protein 2)-like protein (Gene ID: *FUN_017843*; Gundappa et al. 2022). The predicted protein product of this gene contains an AAA domain, hence can bind and hydrolyse ATP (Lupas and Martin 2002). Proteins with these domains have been shown to be involved in several mechanical cell processes, including protein folding. Further characterization of this *N4BP2*-like protein would help better understand the genetic component of growth variation in oysters. Nevertheless, considering that the candidate allele on *N4BP2* explained a small percentage of the phenotypic variance, independent oyster populations should first be evaluated to confirm the validity of the association signal. A second genome-wide significant association – detected only in the TW GWAS – was located in the exon of an uncharacterized gene (*FUN_018833*) whose product shares a high sequence identity (>90%) with similarly uncharacterized proteins in *C. gigas* and *C. virginica* (NCBI accession numbers XP_011433755 and XP_022325737, respectively). Additional genes within the two genomic regions (+/- 100 kb) showing significant associations with flat oyster growth traits are shown in Table S2. Given that the SNPs identified in this study had a small effect on the traits in question, GS would be an effective approach for increasing genetic gains from selection.

### 3.3 Genomic selection

The incorporation of genetic markers into breeding programmes requires a previous understanding of the genetic architecture of the targeted trait(s). In the *O. edulis* population under study, the genetic contribution to the observed variation in growth-related traits was largely polygenic in nature. For the improvement of polygenic traits, genomic selection has been shown to be superior to alternative marker-aided selection due to genome-wide markers capturing a higher proportion of the genetic variation in a trait compared to individual QTL-targeted markers. Consequently, by means of applying GS, higher predictions have been achieved for several production traits in a wide range of commercially important aquaculture species (reviewed in Houston et al. (2020)). Despite GS not yet being widely operational in oyster breeding programmes (Boudry et al. 2021), studies have demonstrated the potential of incorporating genome-wide information into selection schemes in these taxa. In the Pacific oyster, Gutierrez et al. (2018) showed that prediction accuracies for growth-traits increased 25-30% when the genetic merit of individuals was estimated from SNP markers using the Genomic Best Linear Unbiased Prediction (GBLUP) model (VanRaden 2008) compared to a classical pedigree-based approach (PBLUP). Similar results were reported in the Portuguese oyster, as prediction accuracies increased 7-42% for growth-related traits when EBVs obtained by GBLUP were compared to those obtained by PBLUP (Vu et al. 2021a). Since the flat oyster population under study derived from a mass-spawning event, the pedigree structure was unknown. Therefore, comparisons between pedigree and genome based methods for estimating breeding values (e.g. GBLUP and Bayesian approaches) could not be performed.

One of the major barriers of implementing GS is the high number of markers required to accurately predict EBVs and the cost of genotyping these markers (Goddard and Hayes 2007). Therefore, the design of a strategy to reduce the cost of genotyping is critical for the extensive adoption of genomic prediction approaches in aquaculture breeding programmes. One such strategy involves genotyping the minimum number of markers required to achieve maximal accuracy, which by definition is equal to that obtained with a full panel of markers. As shown by Kriaridou et al. (2020) for different aquaculture species, the use of low-density SNP panels has the potential to achieve similar EBV accuracies as when using medium density genotype datasets of around 7-14K SNPs. The authors estimated that only 1,000 to 2,000 SNPs are required to achieve maximal accuracy. These results were shown to be consistent across a range of traits (e.g. disease resistance, growth) and species (e.g. Pacific oyster, Atlantic salmon) showing robustness to differences in family structure, genotyping approach, trait heritability and the underlying genetic architecture. In agreement with these findings, maximal accuracy was attained herein for all the assessed growth-related traits at a minimum density of 2K SNPs, with only a slight decline in accuracy observed at the lower densities evaluated (Figure 4A). Consequently, a reduction in the costs of applying GS for improving growth traits in *O. edulis* can be achieved by means of exploiting low-density SNP panels. Although low-density panels might not accurately capture the genetic resemblance among individuals within a population, and therefore show reduced genetic variance estimations and EBV predictions when compared with high density panels, their use has been widely evaluated and suggested for different aquaculture species and traits. Furthermore, studies have shown that low-density panels can achieve higher accuracies than the classical pedigree-based approach, being a feasible alternative to identify candidates with the highest genetic merit. For example, in rainbow trout (*Oncorhynchus mykiss*) Vallejo et al. (2018) showed that at least 200 SNPs could exceed PBLUP accuracies for bacterial cold water disease resistance. Whilst Al-Tobasei et al. (2021) found a similar trend when using between 500-1000 SNPs for fillet yield traits. To date, the utilization of low-density panels to decrease the cost of genomic evaluations has also been tested in several aquaculture species, including Atlantic salmon (*Salmo salar*) (Correa et al. 2017; Tsai et al. 2016), rainbow trout (Yoshida et al. 2018; Al-Tobasei et al. 2021), and Nile tilapia (*Oreochromis niloticus*) (Barría et al. 2021; Yoshida et al. 2019), suggesting that the development of cost-effective strategies for applying GS will be key for shaping modern aquaculture breeding programs.

**Figure 4.**
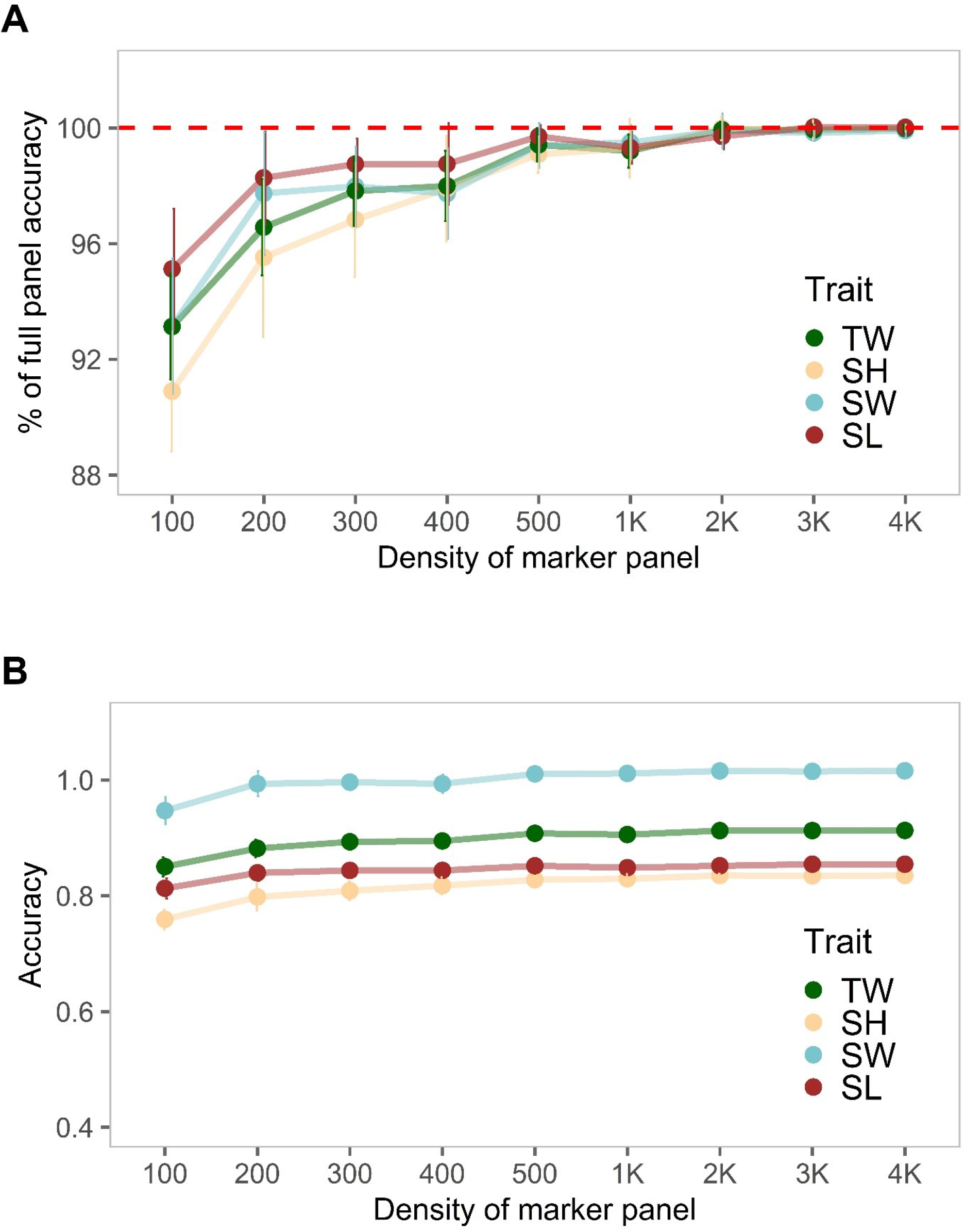
Evaluation of the effect of SNP density on genomic predictions of growth related traits in a European flat oyster population. (A) Percentage of the maximum genomic prediction accuracy achieved using different lower density SNP panels. Values were calculated by dividing the mean accuracy (averaged over ten replicates) estimated at each nominal SNP density by the accuracy obtained using the full SNP dataset. (B) Average genomic prediction accuracy values obtained for oyster growth traits at different panel densities.

Although our results highlight the possibility of reducing the genotyping costs associated with genomic prediction approaches, caution should be taken as even for the smallest marker density (i.e. 100 SNPs), prediction accuracies (averaged over 10 replicates) were high and close to the value obtained with the full marker panel. By using only 100 SNPs the estimated decrease in the accuracy of genomic breeding values (GEBVs) was of 5% for SL, 7% for TW and SW, and 10% for SH (Figure 4A). These values highly exceed those reported in the literature for aquaculture species, where reductions >20% were estimated for panels with 100 SNPs compared to a complete dataset (Kriaridou et al. 2020). The relative stability of GEBVs observed across different marker densities (Figure 4B) is likely explained by the underlying genetic structure of the dataset. For this study, 40 potential parents were placed in the same tank and spawned during successive events. The genetic analysis of the progeny revealed that the population was dominated (70% of the sample size; n=589) by a group of highly related individuals (Figures S2-S3), suggesting there was a large variance in reproductive success among breeders, as previously reported in mass spawning of oysters (Lallias et al. 2010). In the context of GS, the inclusion of highly related animals in the training and validation sets results in only a small number of markers being required to capture the haplotype effects, as related animals share longer haplotypes (Hickey et al. 2014). The fact that in the current study animals grouped in the reference and validation data sets were highly related would have likely increased the accuracy of predicted gEBV, even when animals are genotyped at low density, as also shown by Fraslin et al. (2022) in Atlantic salmon. Moreover, since the high accuracies predicted in the flat oyster population could have also been affected by factors such as a low effective population size (Lee et al., 2017) and the observed structure (Werner et al. 2020), additional populations with different genetic background should be assessed to obtain estimates to be expected in practical breeding programs. Future work focused on evaluating the extent to which low-density panels and alternative strategies (e.g. genotype imputation) can be used to reduce genotyping costs will be key for the cost-effective exploitation of GS by oyster breeding programmes.

## 4. Conclusion

Growth-related traits in *O. edulis* had moderate-low heritability estimates, ranging from 0.22 (for SW) to 0.45 (for TW). High genetic correlations were identified between all traits (>0.9); hence, TW - a trait easier to measure - can potentially be used as a proxy phenotype for improving the three examined morphometric traits (SH, SW and SL). The GWAS results revealed that growth traits were largely polygenic, but with two distinct QTLs on chromosome 4 reaching genome-wide significance. Prediction accuracies were high for all traits (>0.83), with minimal differences observed when comparing estimates obtained using different marker densities. Altogether, these results suggest that the high prediction accuracies found in this study could have been influenced by the uneven family structure of the experimental population. Although low-density SNP panels appear as a promising cost-effective GS strategy, additional populations with different degrees of genetic relationship should be assessed to derive estimates of prediction accuracies to be expected in practical breeding programmes in oysters.

## 5. Acknowledgments

The authors acknowledge funding from the Biotechnology and Biological Sciences Research Council (BBSRC), including Institute Strategic Programme grants (BBS/E/D/20002172, BBS/E/D/30002275 and BBS/E/D/10002070), a grant within the AquaLeap project (BB/S004181/1), funding from Blue Marine Foundation and National Fish and Wildlife Foundation (NFWF). The authors would also like acknowledge MDL Port Hamble Marina for allowing positioning of oyster cages within the Marina, and thank Eric Harris-Scott, Matthew Sanders, Monica Fabra, Tim Regan and Zenaba Khatir for their invaluable help with setup and sampling of the field experiment.

