## Supplemental tables and figures for "Genome-wide association and genomic prediction of growth traits in the European flat oyster (*Ostrea edulis*)"

### Additional figures and tables

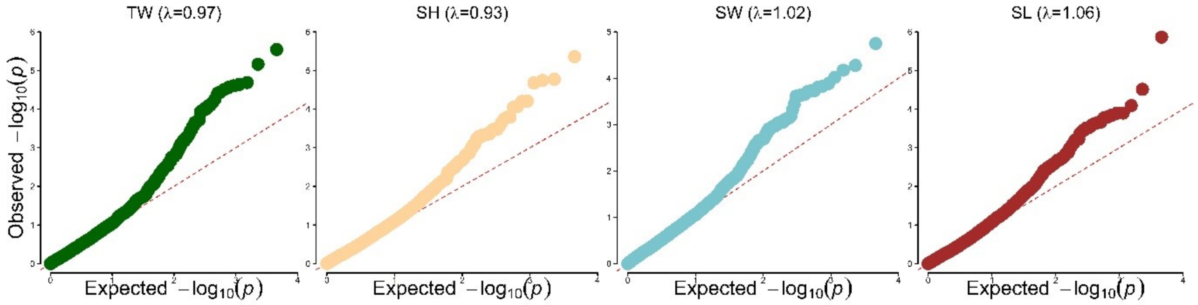


**Figure S1:** Quantile-quantile (Q-Q) plots showing the distribution of the expected (red dashed line) versus observed *P*-values of the GWAS of growth-related traits.

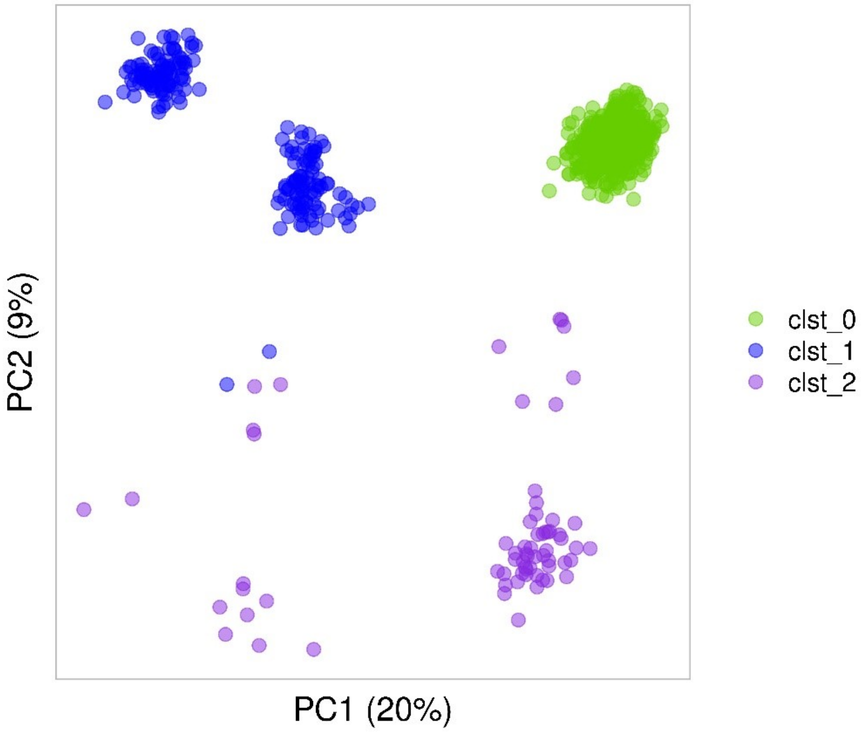


**Figure S2:** PCA of the *O.edulis* population under study showing the dominance of single cluster comprised of highly related individuals (clst_0)

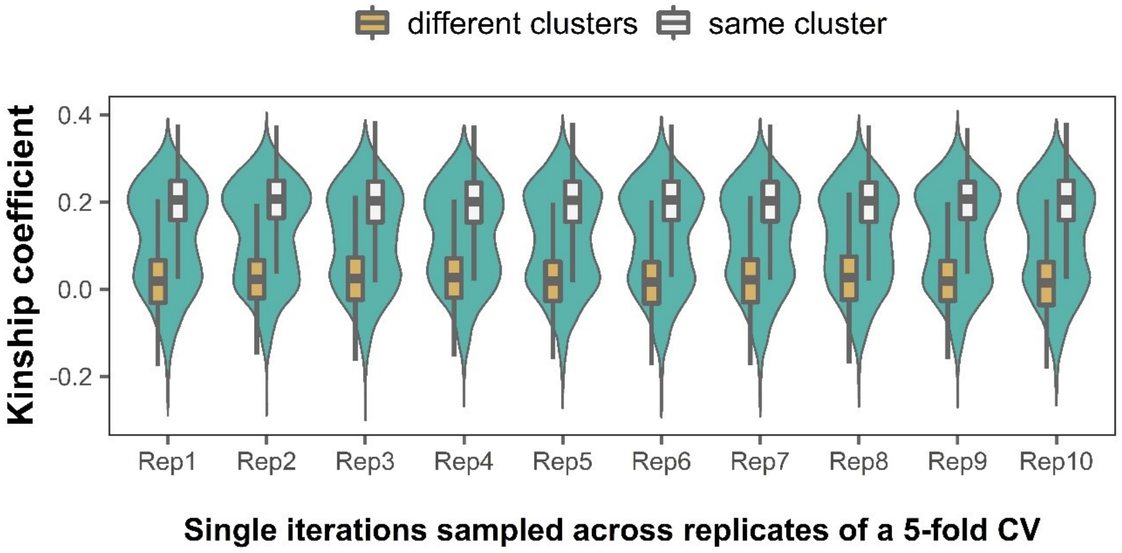


**Figure S3:** Example of the pairwise kinship coefficient between individuals in the training and validation set across randomly selected iterations of different 5-fold cross validation (CV) replicates (10 replicates in total). Boxplots show the distribution of values for pairs of individuals (one from the training set and one from the validation set) belonging to the same or different population clusters. Relatedness was inferred using the KING-robust method implemented in Plink v2.0.

**Table S1:** Details of the SNPs showing a significant (**) and suggestive association with TW

| | | | | | | HWE *P*-value | | | |
|---|---|---|---|---|---|---|---|---|---|
| **Chr** | **Position** | **SNP ID** | **A1** | **A0** | **LRT *P*-value** | **All** | **cluster1** | **cluster2** | **cluster3** |
| 4 | 43233612 | AX-169174635 | G | A | **2.9E-06****| 3.5E-26 | 3.3E-33 | 4.7E-04 | 1.5E-01 |
| 4 | 63862900 | AX-169166246 | T | C | **6.9E-06**** | 7.0E-12 | 3.6E-15 | 1 | 1.9E-01 |
| 4 | 68332666 | AX-169182460 | C | T | 2.1E-05 | 1.2E-11 | 1.3E-14 | 1 | 1.9E-01 |
| 4 | 63376888 | AX-169179861 | A | G | 2.3E-05 | 7.0E-12 | 6.8E-15 | 1 | 1.9E-01 |
| 4 | 59634534 | AX-169201070 | C | T | 2.5E-05 | 1.2E-11 | 6.5E-15 | 1 | 1.9E-01 |
| 4 | 42087933 | AX-169187958 | A | G | 2.7E-05 | 7.8E-19 | 1.5E-33 | 1 | 2.2E-01 |
| 1 | 61517695 | AX-169160998 | A | G | 3.1E-05 | 4.2E-43 | 4.1E-65 | 1 | 4.0E-02 |
| 4 | 41993042 | AX-169195918 | G | A | 3.6E-05 | 3.8E-11 | 4.8E-14 | 1 | 1.1E-01 |
| 4 | 51325140 | AX-169159905 | G | A | 3.9E-05 | 3.2E-27 | 3.5E-33 | 6.0E-01 | 1.6E-01 |
| 4 | 63124249 | AX-169172980 | G | A | 5.6E-05 | 2.5E-16 | 3.4E-15 | 1.4E-01 | 1.1E-01 |
| 4 | 68684016 | AX-169195406 | T | G | 7.0E-05 | 2.4E-24 | 4.6E-32 | 1 | 1.4E-02 |
| 4 | 42973171 | AX-169179093 | T | C | 8.1E-05 | 7.4E-23 | 3.3E-33 | 1 | 1.6E-01 |
| 1 | 64314592 | AX-169175717 | C | T | 8.5E-05 | 1.2E-12 | 2.0E-04 | 4.9E-13 | 1.9E-01 |
| 1 | 64407681 | AX-169199161 | G | T | 1.0E-04 | 4.1E-04 | 2.0E-04 | 1 | 3.4E-01 |
| 1 | 73944671 | AX-169164517 | G | A | 1.0E-04 | 4.1E-04 | 2.0E-04 | 1 | 3.4E-01 |

**Table S2:** Candidate genes in the regions (+/- 100kb) surrounding the significant GWAS hits for TW. Genes with a functional annotation are underlined.

| Chr | Position | SNP ID | Candidate genes |
|---|---|---|---|
| 4 | 43233612 | AX-169174635 | FUN_017842, FUN_045437, **N4BP2_1**, **N4BP2_2**, FUN_017845, FUN_017846 |
| 4 | 63862900 | AX-169166246 | FUN_018829, **SRSF6**, FUN_018831, FUN_018832, FUN_018833, FUN_018834, FUN_018835, **UCK2**, **SURF1**, FUN_018838 |
